# Substrate transport limits phenylalanine ammonia-lyase activity in engineered *Lacticaseibacillus rhamnosus* GG

**DOI:** 10.64898/2026.03.19.713057

**Authors:** Debika Choudhury, Zachary J. S. Mays, Nikhil U. Nair

## Abstract

Probiotic-based encapsulation offers unique advantages over purified enzymes, such as increased protection from thermal-, pH-, and protease-mediated degradation, for oral therapeutic delivery applications. However, one of the major disadvantages of whole-cell systems is lower reaction rate due to substrate-product transport limitations imposed by the cell membrane and/or wall. In this work, we explore the potential of different lactic acid bacteria (LAB) – *Lacticaseibacillus rhamnosus GG* (LGG), *Lactococcus lactis* (Ll), and *Lactiplantibacillus plantarum* (Lp) – as expression hosts for recombinant *Anabaena variabilis* phenylalanine ammonia-lyase (*Av*PAL^*^). *Av*PAL^*^ is used as a therapeutic to treat Phenylketonuria (PKU), a rare autosomal recessive metabolic disorder. Among the three species tested, LGG showed the highest PAL activity followed by *L. lactis*. Next, we attempted to overcome mass transfer limitation in whole-cell biocatalysts in two ways – expression of heterologous transporters and treatment with different chemical surfactants. Engineered strains expressing heterologous transporters exhibited approximately 3-4-fold increased PAL activity, while chemical treatment did not improve reaction rates. This work highlights the challenges and advances in realizing the potential of LAB as biotherapeutics.

**Impact Statement:** Oral delivery of phenylalanine ammonia-lyase (PAL) using engineered probiotics is a promising therapeutic strategy to treat Phenylketonuria (PKU). Although PAL expression has been reported in probiotic strains of *Limosilactobacillus reuteri, Lactococcus lactis*, and *E. coli*, a systematic comparison of lactic acid bacteria (LAB) is underexplored. This study explores the potential of multiple LAB as hosts for PAL expression and investigates strategies to improve whole cell enzymatic activity. The findings from this study provide a foundation for implementing LAB-based delivery of PAL and indicate an important step towards development of probiotic platform for PKU management.

## Introduction

Phenylketonuria (PKU) is a rare inherited metabolic disorder characterized by the accumulation of phenylalanine (phe) in the body and affects about 1 in 15,000 newborns in the United States [1,2]. It is characterized by deficiency in phenylalanine hydroxylase (PAH) activity, resulting in reduced decomposition of phe to tyrosine (tyr) [3]. If intervention is not swift, phe accumulation can lead to intellectual disabilities, neurological defects, failure to thrive, and in rare cases, death [4].

The most effective treatment strategy for management of PKU includes lifelong restriction of phe in the diet [5]. However, because phe is an essential amino acid, eliminating it completely is challenging. Patients often rely on low phe containing synthetic foods and supplements which are costly [6]. These limitations necessitate complementary practical and affordable treatment strategies that aim to lower systemic phe levels in patients suffering from PKU.

Phenylalanine ammonia lyase (PAL) provides an alternative enzymatic route for phe depletion by converting it into *trans*-cinnamic acid (*t*CA) and ammonium (**Figure 1A**), thereby replacing PAH activity [7-9]. Pegvaliase (Palynziq®), a PEGylated recombinant PAL derived from the cyanobacterium *Anabaena variabilis* (*Av*) is currently the only FDA-approved enzyme substitution therapy for PKU [10,11]. However, its administration via daily subcutaneous injections poses risk for anaphylaxis and other adverse immune-mediated reactions.

**Figure 1.**
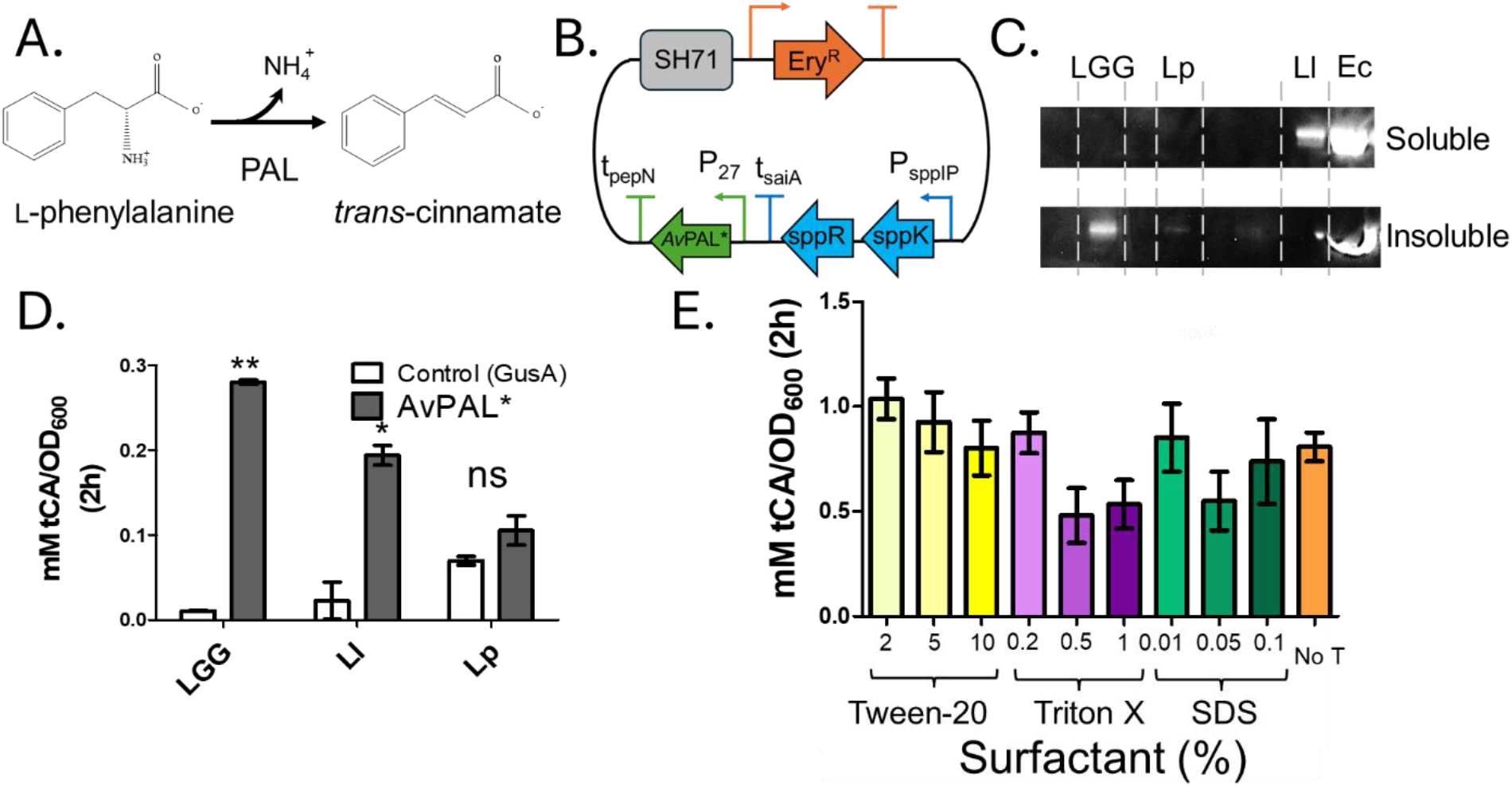
*Av*PAL* expression in LAB. **A)** PAL catalyzes the deamination of L-phenylalanine (phe) to *trans*-cinnamate (*t*CA) with the release of ammonium. **B)** Plasmid map showing the broad host-range plasmid used for *Av*PAL* expression. **C)** Western blot showing recombinant *Av*PAL* in different bacteria. **D)** *Av*PAL* activity in LGG, *L. lactis*, and *L. plantarum*. Control strains were transformed with GusA for comparison. Data represents means from two biological replicates (n=2). ** = *p* < 0.001, ^*^ = *p* < 0.05, ns = p > 0.05 (Students *t*-test). **E)** *Av*PAL* expressing LGG treated different concentration of with surfactants – Tween 20, Triton X-100, and SDS show no significant difference in enzymatic activtiy compared to untreated control. Data represents three biological replicates n = 3, p > 0.05 (One-way ANOVA). LGG = *L. rhamnosus* GG; Lp = *L. plantarum* WCFS1; Ll = *L. lactis* MG1363; Ec = *E. coli* MG1655 *rph*^*+*^. **Alt Text**. Five-panel figures (A-E) showing PAL enzymatic reaction, organization of the *Av*PAL* expression plasmid, PAL expression by Western Blot, PAL activity by measurement of tCA absorbance, and surfactant treatment of recombinant LAB.

Orally-administered PAL can offer a complementary route to lower blood phe levels from dietary and endogenous sources and avoid complications arising from the subcutaneous administration, including adverse injection-site reaction, headache, etc. [9,10]. Oral therapies involving recombinant expression of PAL in engineered bacterial chassis, particularly in probiotics, are a promising avenue. Probiotics are “generally regarded as safe” (GRAS) due to their presence in the gastrointestinal tract (GI) and widespread use in food industry [12]. They have also exhibited potential as oral delivery vehicles for protein drugs as they provide protection from degradation and digestion [13]. Previous reports have demonstrated PAL mediated phe degradation in rodent PKU models using engineered probiotics such as *Lactococcus lactis* and *Limosilactobacillus reuteri* 100-23C (pHENOMMenal) [14,15]. A more recent report using engineered *E. coli* Nissle 1917 (SYNB1618 and SYNB1934), exhibited reduction of blood phe level in both mice and primate has progressed to the clinical trial stage [16-18]. However detailed investigation of different probiotic bacteria as platforms for recombinant PAL expression is limited. Additionally, studies exploring optimization strategies to improve whole-cell bacterial activity remain unexplored.

In this study, we assessed three different lactic acid bacteria (LAB) – *Lacticaseibacillus rhamnosus GG* (LGG), *Lactococcus lactis* MG1363 (Ll), and *Lactiplantibacillus plantarum* WCFS1 (Lp) as whole-cell biocatalysts for recombinant PAL expression and in vitro phe-to-*t*CA conversion. We expressed *Anabaena variabilis* PAL (*Av*PAL^*^) using a synthetic constitutive expression system and measured enzyme activity. Among the LAB tested, *Lacticaseibacillus rhamnosus* GG (LGG) exhibited highest *Av*PAL^*^ activity, followed by *L. lactis*. Because whole-cell biocatalysts are frequently limited by mass transfer and substrate uptake, which negatively impacts the reaction rate, we tried two distinct strategies to improve enzyme reaction rate [19]. We attempted to improve uptake by surfactant-based cell wall decongestion as well as a transporters overexpression strategy to improve enzyme activity. Transporter co-expression increased whole-cell PAL activity by approximately 3–4-fold, whereas surfactant treatment did not produce a significant improvement under the conditions tested.

## Materials and Methods

### Reagents and Enzymes

All cloning enzymes and reagents were purchased from New England Biolabs (Beverly, MA). Oligonucleotide primers were ordered from Eurofins Genomics LLC (Louisville, KY) or GENEWIZ Inc. (Cambridge, MA). Routine molecular biology kits, including plasmid minipreps, PCR purifications, and gel extractions were purchased from Omega Bio-tek (Norcross, GA). For Western blots, mouse-anti-His_6_ antibody (MA1-21315) and goat anti-mouse IgG H & L (AlexaFluor® 488) (A1101) were purchased from ThermoFisher Scientific (Waltham, MA). Media components and chemicals were purchased from Amresco (Solon, OH) or RPI Corp (Mount Prospect, IL).

### Bacterial strains and culture conditions

*Lactiplantibacillus plantarum* WCFS1 (*L. plantarum*, Lp), and *Lacticaseibacillus rhamnosus* GG (LGG), were grown in de Man-Rogosa-Sharpe (MRS) medium. *L. plantarum* was incubated at 30 °C while LGG grew at 37 °C. *Lactococcus lactis* MG1363 (*L. lactis*, Ll) was grown in GM17 medium. All lactic acid bacterial (LAB) cultures were incubated without agitation. *E. coli* NEB5α (Ec) (New England Biolabs, Ipswich, MA) was cultured in Lysogeny Broth (LB) medium at 37 °C with shaking at 225 rpm. For plasmid selection, media was supplemented with erythromycin at concentrations of 5 and 200 μg mL^−1^ for LAB and *E. coli*, respectively. When used simultaneously, erythromycin and chloramphenicol were each added at concentrations 2.5 μg mL^−1^.

### Plasmid and strain construction

Phusion High-Fidelity DNA Polymerase was used for amplifying DNA as per manufacturer’s instructions. Plasmids were propagated in *E. coli* prior to transformation into LAB. Amplified sequences were confirmed by Sanger sequencing. *Av*PAL* and *gusA* were expressed under P_27_ promoter in pSIP411-derived plasmids [20]. For transporter expression, we combined the pAMβ1 origin of replication with the *sppK-sppR* regulatory elements from pJRB01-P_orfX_ plasmid [21,22], the chloramphenicol resistance (Cm^R^) cassette from pNZ8048 plasmid [23], and the synthetic constitutive promoter P_31_ [24], all assembled using the NEB HiFi DNA Assembly kit. All transporter sequences have been included in the Supplementary data. Plasmids were transformed into chemically competent *E. coli* NEB5α following the manufacturer’s instructions. Plasmid DNA was isolated and electroporated into LAB using established protocols [25,26]. Positive transformants were confirmed by colony PCR. For double transformants, plasmids were electroporated sequentially.

### PAL activity assay in whole-cell

Transformants were grown overnight in 5 mL cultures, pelleted, washed with PBS (phosphate-buffered saline), and resuspended in the same to OD_600_ = 5.0. For each assay, 0.5 mL of OD-adjusted cells was pelleted (4,000 × g, 5 min) and the supernatant was replaced with 0.5 mL pre-warmed PBS containing 25 mM L-phenylalanine. Cells were incubated at 37 °C, filtered at 0 h and 2 h, and *t*CA in the filtrate was monitored using absorbance at 290 nm. PAL activity was calculated as a change in A_290_ normalized by the OD_600_ value (**Eqn 1**). This value was converted to mM *t*CA by using a calibration curve, as described previously [27-30].

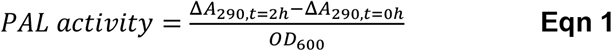

### SDS-PAGE and Western blot

SDS-PAGE and Western blot were performed as described previously [31]. Briefly overnight cultures of PAL transformed LAB were lysed initially in presence of 0.05 % SDS, washed with PBS, followed by lysozyme treatment. Pre-treated cells were lysed on ice in a sonicator. Following sonication, cells were centrifuged at high speed to yield soluble and insoluble fractions. Each fraction was denatured and equal protein concentration from each fraction was loaded into 4–12 % Bis-Tris gels and ran in MOPS (3-(*N*-morpholino)propane sulfonic acid) buffer. For Western blot, the protein bands were transferred to a PVDF (polyvinylidene fluoride) membrane. Following transfer, and blocking, the membrane was incubated sequentially in primary antibody and secondary antibodies. Finally, the signal was developed using Super signal west chemiluminescent substrate.

### Permeabilization assay

Transformed LGG strains from overnight cultures were washed and adjusted to OD 5.0. Next, they were treated with different concentrations of the following surfactants – Tween 20 (2, 5, 10 %), Triton X-100 (0.2, 0.5, 1 %), SDS (sodium dodecyl sulfate) (0.01, 0.05, 0.5%) for 1 hour at 37 °C. Next, permeabilizing agent containing media was removed and cells were washed with PBS and PAL activity was measured as described earlier.

### Statistics and data reproducibility

All data reported here represent biological replicates and were performed on different days. Average data have been plotted with error bars to indicate standard deviation.

## Results and Discussion

### PAL activity is highest in LGG

To directly compare different LAB, we used a broad host range pSH71-based plasmid with a constitutive synthetic P_27_promoter to drive *Anabaena variabilis* PAL (*Av*PAL*) (**Figure 1B**). We confirmed *Av*PAL* expression in all LAB by Western blot and found that the majority of the protein was in the insoluble fractions – and significantly lower expression compared to *E. coli* (**Figure 1C**). Measured OD-normalized PAL activity was highest in LGG, followed by *L. lactis* and *L. plantarum* (**Figure 1D**). Strains expressing *gusA* (encoding *E. coli* β-glucuronidase) served as the negative controls. These results are not altogether surprising since the *Av*PAL^*^ gene sequence used was codon-optimized for *E. coli*, which may explain the low expression level, insolubility, and activity seen for LAB [32,33]. Interestingly, although *L. lactis* showed highest protein expression, it did not translate into higher PAL activity. This could be due to misfolding or improper formation of the MIO (4-methylideneimidazole-5-one) adduct, which is essential for catalysis [34]. Although we tried improving the soluble expression of PAL by growing the LAB at lower temperatures, poor growth led to unreliable OD-normalized activity readings. Since LGG showed highest PAL activity amongst the LAB tested, we used it for all subsequent experiments.

### Permeabilizing agents do not improve whole-cell PAL activity

Previous studies have shown that detergents can enhance biocatalysis through cell wall permeabilization/decongestion. This strategy has been shown to improve whole-cell activity in Gram-positive bacteria for sugar isomerization and epimerization [31,35]. We explored the effect of three different surfactants – Tween 20 (2, 5, 10 %), SDS (0.01, 0.05, and 0.5 %), and Triton X-100 (0.2, 0.5, and 1 %) in overcoming phe transport limitations imposed by encapsulation in LGG cells. Unfortunately, PAL activity did not significantly improve with any of the surfactants (**Figure 1E**). This might indicate that while this strategy worked well for sugar uptake, it may not be effective for amino acids transport into the cell.

### PAL reaction rate improves on transporter co-expression

It was previously reported that expression of high-affinity phe transporter PheP in engineered *E. coli* Nissle 1917 SYNB1618, increased PAL activity [17]. Taking a cue from published work, we tested whether whole-cell PAL activity in LGG could be enhanced by transporter overexpression. Since phe transporters are not well-characterized in most LAB, we used the *E. coli* AroP aromatic amino acid transporter as query to identify putative phe transporters (**Figure 2A**) [36]. Our search revealed candidate phe transporter genes in several Gram-positive bacteria, including those from *Lacticaseibacillus rhamnosus* (WP_014569403.1), *Limosilactobacillus reuteri* (WP_144226493.1), and *Listeria seeligeri* (WP_046328020.1). Apart from these putative transporters, we also expressed the characterized FYWP protein from *Lactococcus lactis* [37]. These transporter genes were expressed from synthetic constitutive promoter, P_31_, in a pAMβ1-based vector (**Figure 2B**). When co-expressed with *Av*PAL in LGG, all transporters increased PAL activity by >2-fold relative to the negative control (**Figure 2C**). This suggests that phe transport limits the PAL activity in LGG even at relatively low expression levels.

**Figure 2.**
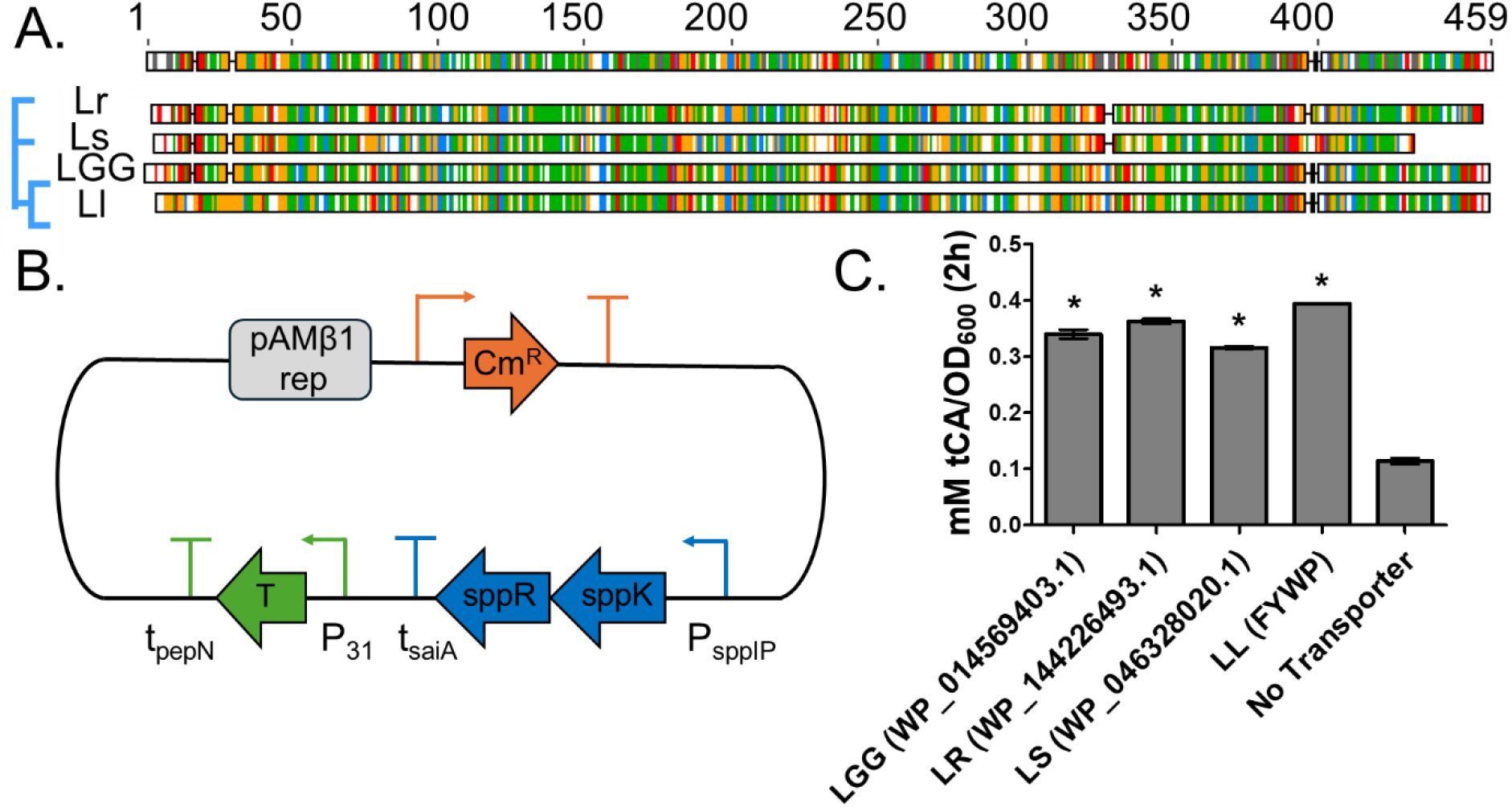
Transporter and PAL co-expression in LGG. **A)** Alignment of amino sequences of the transporters represented by the source organisms using Geneious Prime. All are uncharacterized sequences except for FYWP. Color-coding is based on the physicochemical properties – Green – hydrophobic, blue – basic/positively charged, Red – acidic/negatively charged, Orange/yellow – polar uncharged, White/blank – no residue. The tree at the left indicate sequence similarity-based clustering calculated from the alignment. **B)** Plasmid map showing the broad host-range plasmid used for transporter expression. **C)** *Av*PAL* activity in LGG transformed with transporters from different Gram-positive bacteria (protein accession numbers in parentheses). Data represents means from two biological replicates (n = 2), * = *p* < 0.001 (1-way ANOVA). **Alt Text**. Three-panel figures (A-C) showing alignment of the different transporter sequences from gram-positive bacteria, components of the transporter expression plasdmid, and increase in PAL activity due to transporter co-expression.

## Conclusion

Engineered lactic acid bacteria (LAB) expressing *Av*PAL* represent a promising strategy for phenylketonuria (PKU) management. Here, we compared multiple LAB and identified LGG as the as best hosts for *Av*PAL* expression. Additionally, we investigated approaches to improve whole-cell PAL activity in LGG and found that while heterologous transporter expression enhanced activity, but surfactant treatment had no significant effect. Together, these results serve as a foundation for further optimization of LAB-based PAL delivery systems and indicate the potential of probiotic bacteria as whole-cell platforms for PKU therapy.

## Acknowledgements

This study was supported by NIH grant DP2HD91798 to N.U.N. We would like to thank Dr. Josef R. Bober, Dr. Vikas D. Trivedi, and Dr. Todd C. Chappell for their insightful discussion.

## Data Availability

Sequences for all transporters used in this study are provided in Supplementary Data. All other data will be available from the corresponding author on reasonable request.

## Author Contributions

Debika Choudhury: Methodology, Formal analysis, Investigation, Writing - Original Draft, Visualization, Zachary J. S. Mays: Investigation. Nikhil U. Nair: Conceptualization, Methodology, Writing - Review & Editing, Supervision, Funding acquisition.

## References

1. Stone WL, Los E. Phenylketonuria (PKU). StatPearls [Internet]. StatPearls Publishing; 2023.

2. van Spronsen FJ, Blau N, Harding C, Burlina A, Longo N, Bosch AM. Phenylketonuria. Nature Reviews Disease Primers. 2021/05/20 2021;7(1):36. doi:10.1038/s41572-021-00267-0

3. Elhawary NA, AlJahdali IA, Abumansour IS, et al. Genetic etiology and clinical challenges of phenylketonuria. Human genomics. Jul 19 2022;16(1):22. doi:10.1186/s40246-022-00398-9

4. Rovelli V, Longo N. Phenylketonuria and the brain. Molecular genetics and metabolism. May 2023;139(1):107583. doi:10.1016/j.ymgme.2023.107583

5. Longo N, Hamazaki T, Hollander S, MacDonald A, Muntau AC, Schwartz IVD. Global considerations for lifelong management and therapeutic development for phenylketonuria. Genetics in medicine: official journal of the American College of Medical Genetics. Nov 2025;27(11):101540. doi:10.1016/j.gim.2025.101540

6. Kumar Dalei S, Adlakha N. Food Regime for Phenylketonuria: Presenting Complications and Possible Solutions. Journal of multidisciplinary healthcare. 2022;15:125–136. doi:10.2147/jmdh.S330845

7. Levy HL, Sarkissian CN, Scriver CR. Phenylalanine ammonia lyase (PAL): From discovery to enzyme substitution therapy for phenylketonuria. Molecular genetics and metabolism. Aug 2018;124(4):223–229. doi:10.1016/j.ymgme.2018.06.002

8. Sarkissian CN, Kang TS, Gámez A, Scriver CR, Stevens RC. Evaluation of orally administered PEGylated phenylalanine ammonia lyase in mice for the treatment of Phenylketonuria. Molecular genetics and metabolism. Nov 2011;104(3):249–54. doi:10.1016/j.ymgme.2011.06.016

9. Sarkissian CN, Shao Z, Blain F, et al. A different approach to treatment of phenylketonuria: Phenylalanine degradation with recombinant phenylalanine ammonia lyase. 1999;96(5):2339–2344. doi:10.1073/pnas.96.5.2339

10. Burton BK, Longo N, Vockley J, et al. Pegvaliase for the treatment of phenylketonuria: Results of the phase 2 dose-finding studies with long-term follow-up. Molecular genetics and metabolism. Aug 2020;130(4):239–246. doi:10.1016/j.ymgme.2020.06.006

11. Hydery T, Coppenrath VA. A Comprehensive Review of Pegvaliase, an Enzyme Substitution Therapy for the Treatment of Phenylketonuria. Drug target insights. 2019;13:1177392819857089. doi:10.1177/1177392819857089

12. Pogačar MŠ, Mičetić-Turk D, Fijan S. Chapter 31 -Probiotics: current regulatory aspects of probiotics for use in different disease conditions. In: Dwivedi MK, Amaresan N, Sankaranarayanan A, Kemp EH, eds. Probiotics in the Prevention and Management of Human Diseases. Academic Press; 2022:465–499.

13. Sahoo D, Rodriguez E, Nguyen K, Chintapula U. Probiotic Bacteria as Therapeutics and Biohybrid Drug Carriers: Advances, Design Strategies, and Future Outlook. ACS Applied Bio Materials. 2025/09/15 2025;8(9):7513–7534. doi:10.1021/acsabm.5c00959

14. Durrer KE, Allen MS, Hunt von Herbing I. Genetically engineered probiotic for the treatment of phenylketonuria (PKU); assessment of a novel treatment in vitro and in the PAHenu2 mouse model of PKU. PloS one. 2017;12(5):e0176286. doi:10.1371/journal.pone.0176286

15. Jia X, Liu J, Xiang H. [A new strategy of gene therapy for hyperphenylalaninemia rats]. Zhonghua yi xue za zhi. Jun 2000;80(6):464–7.

16. Charbonneau MR, Denney WS, Horvath NG, et al. Development of a mechanistic model to predict synthetic biotic activity in healthy volunteers and patients with phenylketonuria. Communications biology. Jul 22 2021;4(1):898. doi:10.1038/s42003-021-02183-1

17. Isabella VM, Ha BN, Castillo MJ, et al. Development of a synthetic live bacterial therapeutic for the human metabolic disease phenylketonuria. Nature Biotechnology. 2018/09/01 2018;36(9):857–864. doi:10.1038/nbt.4222

18. Vockley J, Sondheimer N, Puurunen M, et al. Efficacy and safety of a synthetic biotic for treatment of phenylketonuria: a phase 2 clinical trial. Nature metabolism. Oct 2023;5(10):1685–1690. doi:10.1038/s42255-023-00897-6

19. Sakkos JK, Wackett LP, Aksan A. Enhancement of biocatalyst activity and protection against stressors using a microbial exoskeleton. Scientific reports. Feb 28 2019;9(1):3158. doi:10.1038/s41598-019-40113-8

20. Karlskås IL, Maudal K, Axelsson L, Rud I, Eijsink VG, Mathiesen G. Heterologous protein secretion in Lactobacilli with modified pSIP vectors. PloS one. 2014;9(3):e91125. doi:10.1371/journal.pone.0091125

21. Diep DB, Mathiesen G, Eijsink VG, Nes IF. Use of lactobacilli and their pheromone-based regulatory mechanism in gene expression and drug delivery. Current pharmaceutical biotechnology. Jan 2009;10(1):62–73. doi:10.2174/138920109787048571

22. Oh J-H, van Pijkeren J-P. CRISPR–Cas9-assisted recombineering in Lactobacillus reuteri. Nucleic Acids Research. 2014;42(17):e131–e131. doi:10.1093/nar/gku623 %J Nucleic Acids Research

23. Martinez-Jaramillo E, Garza-Morales R, Loera-Arias MJ, et al. Development of Lactococcus lactis encoding fluorescent proteins, GFP, mCherry and iRFP regulated by the nisin-controlled gene expression system. Biotechnic & histochemistry: official publication of the Biological Stain Commission. 2017;92(3):167–174. doi:10.1080/10520295.2017.1289554

24. Rud I, Jensen PR, Naterstad K, Axelsson L. A synthetic promoter library for constitutive gene expression in Lactobacillus plantarum. Microbiology (Reading, England). Apr 2006;152(Pt 4):1011–1019. doi:10.1099/mic.0.28599-0

25. Aukrust TW, Brurberg MB, Nes IF. Transformation of Lactobacillus by Electroporation. In: Nickoloff JA, ed. Electroporation Protocols for Microorganisms. Humana Press; 1995:201–208.

26. Keersmaecker SCJD, Braeken K, Verhoeven TLA, et al. Flow Cytometric Testing of Green Fluorescent Protein-Tagged Lactobacillus rhamnosus GG for Response to Defensins. 2006;72(7):4923–4930. doi:10.1128/AEM.02605-05

27. d’Amone L, Trivedi VD, Nair NU, Omenetto FG. A Silk-Based Platform to Stabilize Phenylalanine Ammonia-lyase for Orally Administered Enzyme Replacement Therapy. Molecular pharmaceutics. Dec 5 2022;19(12):4625–4630. doi:10.1021/acs.molpharmaceut.2c00512

28. Mays ZJ, Mohan K, Trivedi VD, Chappell TC, Nair NU. Directed evolution of Anabaena variabilis phenylalanine ammonia-lyase (PAL) identifies mutants with enhanced activities. Chemical communications (Cambridge, England). May 14 2020;56(39):5255–5258. doi:10.1039/d0cc00783h

29. Trivedi VD, Chappell TC, Krishna NB, et al. In-Depth Sequence–Function Characterization Reveals Multiple Pathways to Enhance Enzymatic Activity. ACS Catalysis. 2022/02/18 2022;12(4):2381–2396. doi:10.1021/acscatal.1c05508

30. Trivedi VD, Mohan K, Chappell TC, Mays ZJS, Nair NU. Cheating the Cheater: Suppressing False-Positive Enrichment during Biosensor-Guided Biocatalyst Engineering. ACS synthetic biology. Jan 21 2022;11(1):420–429. doi:10.1021/acssynbio.1c00506

31. Bober JR, Nair NU. Galactose to tagatose isomerization at moderate temperatures with high conversion and productivity. Nature communications. Oct 7 2019;10(1):4548. doi:10.1038/s41467-019-12497-8

32. Gustafsson C, Govindarajan S, Minshull J. Codon bias and heterologous protein expression. Trends in Biotechnology. 2004/07/01/ 2004;22(7):346–353. doi: 10.1016/j.tibtech.2004.04.006

33. Johnston C, Douarre PE, Soulimane T, et al. Codon optimisation to improve expression of a Mycobacterium avium ssp. paratuberculosis-specific membrane-associated antigen by Lactobacillus salivarius. Pathogens and Disease. 2013;68(1):27–38. doi:10.1111/2049-632X.12040 %J Pathogens and Disease

34. Schwede TF, Rétey J, Schulz GE. Crystal Structure of Histidine Ammonia-Lyase Revealing a Novel Polypeptide Modification as the Catalytic Electrophile. Biochemistry. 1999/04/01 1999;38(17):5355–5361. doi:10.1021/bi982929q

35. Park C-S, Kim T, Hong S-H, Shin K-C, Kim K-R, Oh DKJPO. D-Allulose Production from D-Fructose by Permeabilized Recombinant Cells of Corynebacterium glutamicum Cells Expressing D-Allulose 3-Epimerase Flavonifractor plautii. 2016;11

36. Chye ML, Guest JR, Pittard J. Cloning of the aroP gene and identification of its product in Escherichia coli K-12. Journal of bacteriology. Aug 1986;167(2):749–53. doi:10.1128/jb.167.2.749-753.1986

37. Trip H, Mulder NL, Lolkema JS. Cloning, Expression, and Functional Characterization of Secondary Amino Acid Transporters of Lactococcus lactis. 2013;195(2):340–350. doi:10.1128/jb.01948-12

