## Supplemental Information for "Substrate transport limits phenylalanine ammonia-lyase activity in engineered *Lacticaseibacillus rhamnosus* GG"

Debika Choudhury^1^, [Zachary J. S. Mays](https://pubmed.ncbi.nlm.nih.gov/?term=%22Mays%20ZJS%22%5bAuthor%5d)^1^, Nikhil U. Nair^1,*^

^1^ Department of Chemical & Biological Engineering, Tufts University, Medford, MA 02155

**Transporter sequences**

1. *Lacticaseibacillus rhamnosus* GG permease

>source=Lacticaseibacillus_rhamnosus_GG|ref=RefSeq:WP_014569403.1|family=AnsP|COG=COG1113|product=amino_acid_permease|this_study_CDS

ATGCAAGCCGAAAAAGATGATCGGCCTGAACTAGCACGTCATTTGAAGAGTCGCCATGTTCAACTGATTGCAATAGGCGGCACGATTGGGACTGGCTTGTTTTTGGGCTCTGGTCAATCGATTCACTCGGCAGGACCGTCCATTTTATTTGCTTATTTAATAACTGGTGGCATTTGCTTTTTGCTAATGCGTGCATTAGGCGAGTTGTTGATGTCTGATGTGGATTCACATACTTTTATTGATTTCATCACAAAATACCTGGGTATGGATGCCGGTTGGGTGACTGGCTGGACTTACTGGATATGCTGGGTAGCACTTGCGATGGCTGAAGTGACGGCGATTGGATTATATATTCGGTTCTGGTTGCCTGATGTGCCGCAGTGGCTACCCGGTTTAATTGCTTTAGCGATCTTGTTGTTGTTGAATCTGGTCAGTGTCGGATTATTCGGAGAAGCAGAATTCTGGTTTGCCTTGATTAAAATTGTGGCCATCATTGGACTCATCGTTTTAGGTATCTTCATGGTGATTGTTCGCTTTAAAACGCCACTTGGTCATGCCAGTCTTGGCAATCTTGTTAACGATGGCGGCTTTTTCCCTAAAGGCGTTGGCGGTTTTCTCATGTCACTGCAGATGGTTGTGTTTAGTTTTGTCGGCATTGAAATGGTTGGCTTGACTGCTAGTGAGACCAAAAATCCACATAAGGTGATTCCGGAAGCGATCAACGAAATTCCAATGCGGATTTTGCTGTTTTATGTGGGGGCCTTGTTTGTCATCATGTGTATTTATCCATGGCGGCATGTCTCCCCGGTTAACAGCCCGTTTGTTGAAGTCTTTAATAATGTGGGGATTCCGTTTGCGGCCGATATCATTAACTTTGTGGTATTAACCGCGGCGGCCTCGGCTTGTAATTCATCTATTTTTAGCACGGGGCGCTTGTTATTTTCGCTAACGTTGAACGGAAAAAGTCAATCAGCGCAGTGGACTGCCAAACTTTCACGACGCCAGGTGCCAGCTCGGGCGATTCTGGTCTCAACCGGTGTCATTGCCGTAGCGGTTGTCTTGAACCTTTTTCTTCCCGGTTCGGTTTTCACGCTGGTTTCAAGTATGGCAACGATCAGTTTTCTGTTTGTTTGGGGCATGATCATGTTGGCACATTTGCGGTATAAGAAGTTGCATCCCCACAGTACCGACTTTCGGATGCCATTGTACCCATTTGCTGATTATCTTGTGCTGGGTTTTCTGCTATTAACCGCAGTAATCATGATGTTTGATCGTGCAATGTTATCGGCGCTGATCTTCGCAATTGTGTGGATTGCCACACTCTTTATATTGCGGCGGCTGCGGCGAGCGGAAAAAGCAGCGTAA

2. *Limosilactobacillus reuteri* permease

>source=Limosilactobacillus_reuteri|ref=RefSeq:WP_144226493.1|family=AnsP|COG=COG1113|product=amino_acid_permease|this_study_CDS

ATGGACGAAAAACAACAACAGAAGTTAGAACGATCGCTTAAAAGTCGTCACGTGACGATGATTGCTATTGGTGGGGCTATTGGGACCGGTCTTTTTCTTGGATCAGGAACAGCTATTCACCAAGCAGGTCCATCAATCATTCTTTCATACCTGATCGTCGGTATCTTTTGCTTCTTTATGATGCGGGCGTTAGGGGAACTTATTTTAGCTGATACTTCAAAACATTCATTTATTGATTCGGTTAAGGAATACCTTGGTGACCGAATGGAATTTGTTGCCGGCTGGATGTACTGGGCATGTTGGCTGACATTAGCAATGGCTGACCTTACTGCAACTGGTATTTACTTGAAATATTGGTTTCCGAATTTACCACAATGGGTTGGGCCGTTAATTATTGTTATTTTATTAATGTTAGTAAACATGGTAAACGTTGGGCTGTTTGGTGAACTAGAAAGTTGGTTCTCCATGATTAAGGTGATTGCAATTTTAGCCTTAATAGCAGTTGGCGCGGTCTTACTTGTGATGCATGGTCATGTTGAAGGACGGCCGGTAACTCTTTCAAACCTAGTTAACCAGGGTGGCTTTTTCCCTACTGGTCCGATGGGCTTCTTAATGTCTTTCCAAATGGTAGTCTTTGCCTTTGTGGGAATTGAAATGGTAGGATTAACAGCCGGAGAAACTAAGAATCCAGATAAAGATATTCCTAAGGCGATCAATACCTTACCAGTCCGGATTGGTTTGTTCTATATCGGATCAATGATTGCGATGATGTCAATTTACCCATGGTTCCAGATCAAGACTACTTCTAGTCCATTTGTTCAGGTGTTCGCAGCAATCGGTGTTCCAGGAGCAGCTGCGATTTTAAACTTTGTGGTTTTGACGGCTGCAATGTCTGCAACAAACAGTGCGATCTTTAGTACTAGTCGTTCCCTCTACTCATTAGCTCGAAGCGGGAACGCACCAAAACGTTTTGGTGAATTAAGCGCGAAGGCAGTGCCTAATCATGCATTAACTTTCTCTTCATTGATTCTTTTTATCACAGTTATCTTGAATTACATTATGCCAGCAGGAATATTCGATGTAATTGCTGGAATTTCAACAATTACCTTTATTTTTACGTGGATCATTATTTTGGTTGCACATATTAAGTTCCGGCGGCAGAATCCAAAGGGAGTTACGAATTTCAGAATGCCAGGATATCCGATTACTAGTTGGTTAACAATTATTTTCTTCTTGGCAGTTCTAGTAATTTTACTCTTTATTGATTCAACACGAGTTCCATTAATCCTTTCAATCGTTATTTTTGCATTGCTTGCTTATGGTTATGGATTTTTGAAGAAGAAAAATTAA

3. *Listeria seeligeri* permease

>source=Listeria_seeligeri|ref=RefSeq:WP_046328020.1|family=AnsP|COG=COG1113|product=amino_acid_permease|this_study_CDS

ATGGAAGAGCAGAAAGAAGAACTGCAGAGAGGACTGAAAAATAGACATATTCAGTTGATTGCAATAGGTGGAGCAATTGGTACGGGGCTATTTCTAGGAGCCGGAAAGTCAATTCACTTAGCTGGCCCATCTATCATGTTAGTTTACTTAATTATTGGCGCTATTTTATTTTTTGTAATGAGGGCTTTAGGTGAATTGTTAATACATAACCCAACAACAGGATCATTTACAGAATTTGCCGAACAATATATTGGACCGTGGGCAGGCTTTATTACAGGCTGGACATATTGGTTCTGCTGGATTGTCACAGGGATTGCTGAAATAACAGCGGTTGGAATGTACGTTAAATTTTGGGTGCCAGACTTGCCGCAGTGGATACCAGCACTGGGTTGCGTGTTAATTTTACTTCTATTTAATTTAGCGACTGTAAAAGCATTTGGAGAAATTGAATTCTGGTTTGCGATTATCAAAGTAGTCGTAATTATTGCACTCATTGTTATTGGTTTTGTACTTGTATTTATTGGCTACAAACATGGAACAAGTACTGCATCATTTTCTAATTTAGTGGACTATGGCGGCTTTTTCCCAAATGGAGTAGCAGGTTTCTTACTCGCATTCCAAATGGCGACTTTCTCGTTTGTTGGAATTGAGCTTGTTGGGGTGACAGCGGGCGAGGCAGATGATCCAGAACGTACTTTGCCAAAAGCCATTAACAATATTCCTATCCGGATTTTGATATTTTATATTGGTGCTTTACTAGTTTTAATGTCCATTTATCCATGGAGCAATATTGATCCTAATACTAGCCCATTTGTTAGTGTATTTACGATGATAGGAATTCCAGCCGCAGCCGGGATTATTAACTTTGTTGTACTCACAGCTGCAATGTCTTCTTGTAATAGTGGGATTTTTAGCACGAGTCGGATGCTTTACACACTTTCAGCAGAAGGAAAAGCACCGAAAAAAATGCATCATTTAAGCTCAAACGGTGTACCAGCAACTGCTTTAATCACTTCCACTGCTTGTTTACTTATTGGTGTGTTCTTAAATTATGTTTTACCAGAACAAGTATTTATTTTGGTTACAAGTATCGCAACTATTTGCTTTATCTGGGTTTGGGGCGTCATTTTAGTAGCTCATTTACGTTTCCGTAAGAAACATCCAGAAATAGCTGCCAAAAGTAAATTTAAAATGCCATTATCACCATTGATGAACTGGGTTAGCTTAATTTTCTTTGTGGGATTATTAATTATCCTAGGTTTTGCTGCTGATACGCGAATTGCTCTCTTTGTAACACCATTGTGGTTCCTTATTCTAGCCATTGCATATCAAATTGTAAAAGCTTCCAATAAAAAGTCGGAAATTCGACATATTTAA

4. *Lactococcus lactis* permease

>T4|source=Lactococcus_lactis|ref=UniProt:A2RMP5|gene=fywP|product=aromatic_amino_acid_permease|this_study_CDS

ATGGAAAATTCTACAGGCTTGAAGTCTTCTTTAAAGACTCGCCACATTGTTATGCTCTCACTTGGGGGAGCGATTGGATCAGGACTATTTTTAGGTTCTGGTAAAGTCATCGCACAAGCAGGGCCAAGTGTGCTGCTCTCCTACGTCTTAGCTGGGCTGACCCTTTATGTCGTCATGTATGGTGTTGGAAAAATGGTCATTCATCAAGATGACCATAAAGCTGGGATGGCTGGAGTTGTTGCACCTTTTATTGGAGATCATTGGGCGCATTTTGCAGACTGGGTTTACTGGGCAACTTGGATGGCGGTCCTCATCGCAGAAGAAGCGGGAGTTTCAACTTTCCTTGCTATACTGATTCCTGGCGTCCCACTTTGGGTTTTTGCTTTGGTTGTTGCGGTTCTTGGAACGGCAATTAATCTTTGGTCAGTCAAAGCTTTTGCCGAAACGGAATATTGGTTGGCCTTTATCAAGGTCGCTGTTATCCTACTTTTGATTGCTTTGGGTATTTATTTATTGGTTATCAATGATGCTCATCTTGGTTTTGTTGCCGATAGTGCTCAAAAAGTAACTACGAAAAGTACAGCACCAAGTTTTGCTCCAAATGGTTTTTCAGGATTTTTAACTTCACTATTAGTCGTTATTTTCTCATTTGGTGGTTCTGAATTGGCGGCCATTACAGTTGCTGAAACAGAAAATCCAAAAGTGGCTATTCCGAGAGCTATTCGAGGCGTTTTAATTCGGATTATTAGTTTTTATGTTATTCCAATCTTTTTATTCTTGCATTTATTGCCATGGAGCGAAGTGTCAAATCCAGACGCCGCTAGTCCATTTGCGACAATTTTTGCTCGAGTTGGAATTCCTCACGCAGATAAAATTGTACTTGTAATTATTGTTATTGCTATTTTCTCAGCCGTGAATTCGGCAATTTATGCTACTTCACGTTCACTTTATTCTCGAATTCAAGGTTCTAGCACTTATGTGGGTAAAAAATTAGGTAAATTATCAAAAAATCAGGTGCCAACGAATGCGATTTTAGTTTCTTCATTTGTTCTATTTATCGGAGTTCTTTTATCAGCTGTTTTGGGTGATGGTTTCTGGCAATTTGTCGCTGGTTCAATCTCATTTACCATTTCAATCGTATGGATTCTCTTATTAGTTGCTGCTTTAGTTTTGTATTTTAAACATAAAGAAGTCACAAATTGGTTTGTTAAATTGGCAACACTTGTAGTACTTATCGCTTTAAGCCTCGTGTTTATCATGCAAATTATCACAAATCCTTGGACATTGTCCGTCTTTGCCTTGGTTATTTGCCTTTTGTCTTACTTTAGTTACCGCAAAAAAAAAAGCATAATAGAAAAAACATTCATTTTATAAT
